# Change of Guards: ELAVL Proteins Switch miRNA Export Responsibility to Regulate Neuronal Differentiation

**DOI:** 10.1101/2025.08.08.669247

**Authors:** Sritama Ray, Suvendra N Bhattacharyya, Kamalika Mukherjee

## Abstract

ELAVL1 protein HuR controls the stability and activity of miRNAs by disconnecting miRNPs from their target mRNAs and exporting Ago2-uncoupled miRNAs to help hepatic and macrophage cells manage stress. In mammalian neurons, the homolog of HuR, ELAVL4 or HuD, is expressed; however, its role in miRNA regulation remains unknown. PC12, a rat pheochromocytoma cell line, is a well-established model for studying neuronal differentiation that resembles sympathetic neurons. In PC12 cells, HuD levels increase during differentiation, which suppresses HuR expression. Thus, HuD overrides HuR’s control of miRNA activity in differentiating cells, promoting differentiation by regulating the activity and levels of specific miRNAs, such as let-7a and miR-125b. When HuD is present, it decreases the activity of these miRNAs by binding to them, thereby facilitating their export and ensuring neuronal differentiation. Overall, there is a shift in miRNA regulation, with HuR being replaced by HuD in controlling miRNA export during neuronal differentiation. This switch in ELAVL-mediated miRNA regulation is essential for the differentiation process.

## Introduction

Post-transcriptional gene regulation is essential, and during differentiation, neuronal cells change gene regulation, necessitating precise control of protein production that is crucial for forming neural networks and synapses (1). RNA-binding proteins (RBPs) regulate mRNA stability, translation, splicing, and degradation, allowing neurons to respond to signals and ensuring proper differentiation (2-4).

HuR and HuD are key RBPs regulating neuronal differentiation (5, 6). They belong to the Hu family, which has RRMs that bind to AU-rich elements (AREs) in the 3′ UTRs of target mRNAs (6), thereby modulating mRNA stability, localization, and translation. HuR is a ubiquitously expressed, nuclear protein that regulates RNA splicing and post-transcriptional events. During stress, HuR translocates to the cytoplasm, stabilizing and enhancing the translation of mRNAs related to growth, proliferation, and stress responses (7). It also relieves the repression of target mRNAs by displacing repressive miRNAs at the 3′ UTR during starvation (8), helping to release stored mRNAs from RNA processing bodies or P-bodies, sites known for RNA storage and degradation, and recruiting them to polysomes (9). Additionally, HuR mediates the exosome or extracellular vesicles (EV)-mediated export of specific miRNAs, such as miR-122, during nutrient deprivation, thereby adjusting cellular miRNA levels and activity in hepatic cells (10).

HuD is more neuron-specific, expressed in differentiated neurons, and stabilizes and enhances mRNA translation related to neuronal development and synaptic function (11). Unlike HuR, which promotes growth, HuD is linked to inhibiting proliferation and promoting cell differentiation. Despite structural similarities, HuR and HuD are considered to have distinct, opposing roles vital for neuronal function (5).

This study demonstrates that HuD levels increase while HuR levels decrease during nerve growth factor-induced differentiation of PC12 cells, a model that mimics the development of sympathetic neurons (12). HuD expression suppresses HuR and promotes PC12 differentiation. In differentiating cells, HuD regulates miRNAs such as let-7a and miR-125b, inhibitors of neuronal differentiation, by sequestering and exporting them out of differentiating cells, thereby reducing their intracellular levels, promoting PC12 differentiation. This data indicates a shift in miRNA regulation during the differentiation of PC12 cells, where HuD replaces HuR as the primary regulator of miRNA export, which is vital for PC12 differentiation.

## Results

### Reciprocal regulation of HuD and HuR expression during NGF-induced differentiation and upon NGF withdrawal in differentiated PC12 Cells

While examining the altered expression of ELAVL proteins during NGF-induced differentiation of PC12 cells, we observed an increase in HuD expression, along with an elevation in Neurofilament-L protein — a well-established marker of neuronal differentiation in PC12 cells — and a decrease in HuR expression (13). Western blot analysis revealed a time-dependent increase in HuD protein, with significant increases observed at 24, 48, and 72 hours of differentiation (**Fig. 1A-C; Fig. S1A**). Interestingly, although HuD protein levels continued to increase, the mRNA encoding HuD did not rise beyond 48 hours (**Fig. 1D**). Conversely, HuR exhibited a consistent decline at both the protein and mRNA levels during differentiation (**Fig. 1B-D**). The high HuD in differentiated PC12 cells was predominantly cytosolic. At the same time, residual HuR remained nuclear in differentiated PC12 cells (**Fig. 1E**). Withdrawal of NGF from differentiated PC12 cells, leading to dedifferentiation, resulted in decreased levels of Neurofilament-L and HuD proteins, along with an increase in HuR expression (**Fig. 1F-G, Fig.S1B**), indicating an inverse expression pattern of HuR and HuD during differentiation and dedifferentiation. The mRNA levels of HuR and HuD also exhibited patterns similar to those of their respective proteins during NGF withdrawal for differentiated PC12 cells (72h of NGF differentiation before NGF withdrawal) (**Fig. 1H**). These findings suggest a tightly regulated interaction between HuR and HuD, which potentially functions as a molecular switch to control neuronal differentiation.

**Figure 1.**
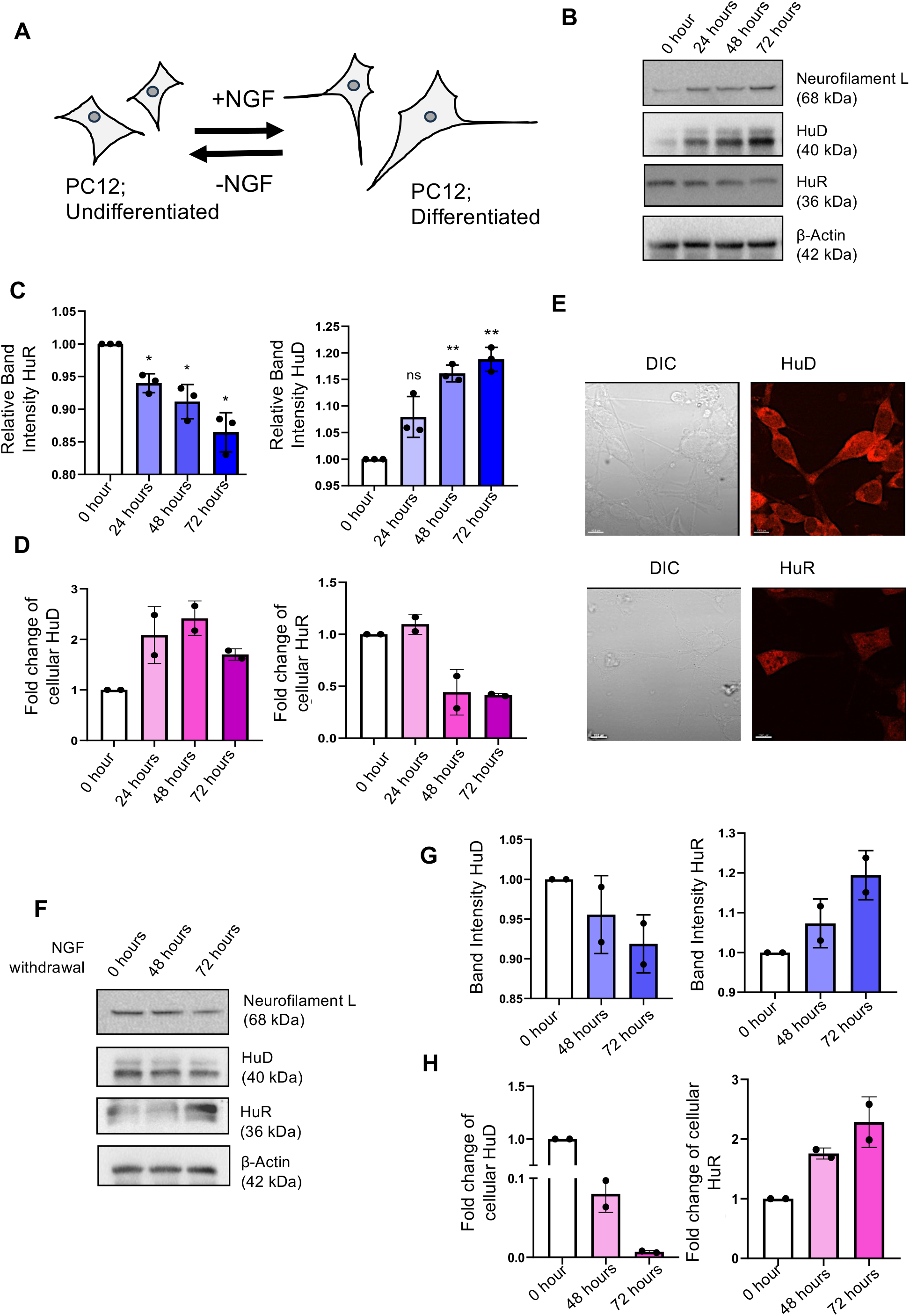
Expression of HuD and HuR protein and mRNAs in differentiating and dedifferentiating PC12 cells. **(A)** Schematic of the experiments. **(B)** Western blot showing the expression levels of Neurofilament-L, HuD, and HuR proteins in PC12 cells differentiated for 0, 24, 48, or 72 hours with 100 ng/mL NGF. β-Actin served as a loading control. **(C)** Relative band intensity of HuR and HuD quantified from three sets of western blots and normalized against β-Actin band intensity. p values 0.0190, 0.0279, 0.0160 for HuR and p values 0.0704, 0.0031, 0.0048; for HuD. **(D)** Relative levels of HuR and HuD mRNAs in PC12 cells differentiated for 0, 24, 48, or 72 hours, quantified by qRT-PCR against GAPDH mRNAs. **(E)** Confocal images displaying HuD (Red) (upper panel) and HuR (Red) (lower panel) expression in 72-hour differentiated PC12 cells. DIC images are also shown. **(F)** Western blot of Neurofilament-L, HuD, and HuR in cells after 0, 48, or 72 hours of dedifferentiation following 72 hours of pre-treatment with 100 ng/mL NGF in medium without NGF. β-Actin was used as a loading control. **(G)** Quantification of HuR and HuD in de-differentiating PC12 cells. Relative quantification was done against β-Actin band intensity for two sets of Western blots. **(H)** Relative levels of HuR and HuD mRNAs in 0, 24, or 72 hours of dedifferentiated PC12 cells, measured by qRT-PCR against GAPDH mRNAs. B-Actin served as a loading control. Data are shown as means ± SDs.

### HuD promotes and HuR restricts neuronal differentiation while inhibiting each other’s expression

HuR is a growth-promoting protein (14), while HuD may be involved in differentiation processes (15). Thus, a reciprocal regulation of both proteins may exist that happens in differentiating neurons. We have used a FLAG-HA-tagged version of the rat HuD protein (FH-HuD) to express it in PC12 cells and examined the role of HuD in NGF-induced differentiation by assessing the expression of key differentiation markers and neurite length in treated cells. FH-HuD expression significantly increased GAP43 and Neurofilament-M mRNA levels and Neurofilament-L protein expression (**Fig. 2A, 2B-D**), suggesting enhanced neuronal differentiation. Additionally, a notable increase in neurite length was observed compared to the control (**Fig. 2C**), indicating that HuD has a positive influence on NGF-induced neuronal differentiation. In contrast, HA-HuR expression resulted in reduced expression of GAP43 and Neurofilament-M at the mRNA level, accompanied by decreased Neurofilament-L protein expression (**Fig. 2E and 2G**). Neurite length was also significantly reduced (**Fig. 2F**), suggesting that HuR, unlike HuD, suppresses neuronal differentiation.

**Figure 2.**
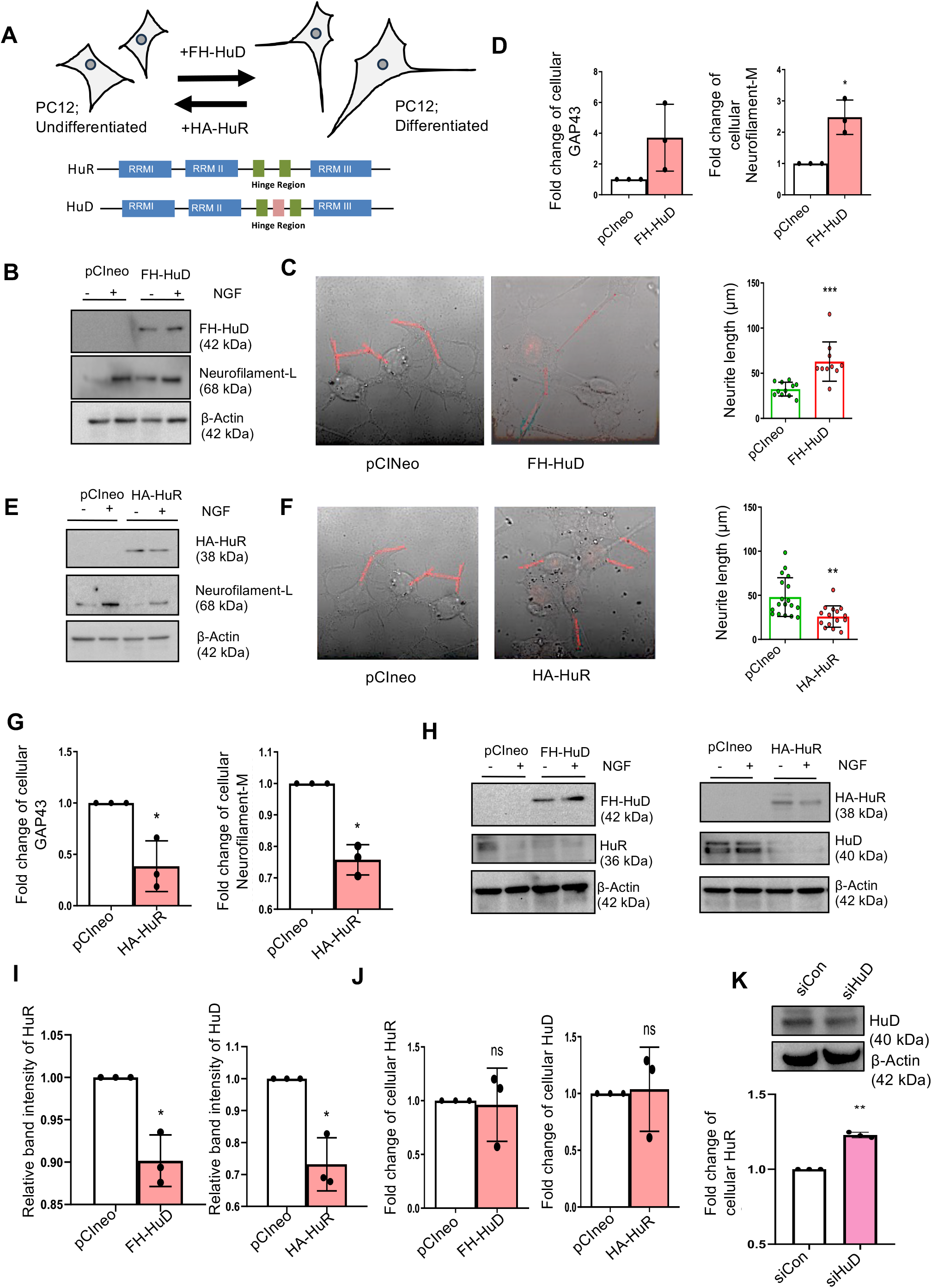
HuD suppresses HuR and augments the differentiation of PC12 cells. **(A)** Scheme of the experiments and HuR and HuD domain structures. **(B)** Western blot images showing FH-HuD and Neurofilament-L proteins in PC12 cells expressing pCIneo or FH-HuD. β-Actin served as a loading control. **(C)** Neurite length measurement in PC12 cells expressing pCIneo or FH-HuD. p = 0.0003; more than 10 cells were measured. **(D)** qRT-PCR quantification of cellular GAP43 and Neurofilament-M mRNA levels, normalized against GAPDH, in PC12 cells expressing pCIneo or FH-HuD. p = 0.0431; n = 3. **(E)** Western blot images depicting HA-HuR and Neurofilament-L protein levels in PC12 cells expressing pCIneo or HA-HuR. β-Actin was used as a loading control. **(F)** Neurite length measurement in PC12 cells expressing pCIneo or HA-HuR. p = 0.0003; over 10 cells measured. **(G)** qRT-PCR quantification of GAP43 and Neurofilament-M mRNA levels, normalized to GAPDH, in PC12 cells expressing pCIneo or HA-HuR. p = 0.0496 (GAP43); p = 0.0129 (Neurofilament-M); n = 3. **(H)** Western blot images showing HA-HuR and Neurofilament-L protein levels in PC12 cells expressing pCIneo, FH-HuD, or HA-HuR. β-Actin served as a loading control. **(I)** Quantification of HuD and HuR protein expression in PC12 cells expressing pCIneo, FH-HuD, or HA-HuR; p = 0.0306, p = 0.0305; n = 3. **(J)** qRT-PCR analysis of HuR (right) and HuD (left) mRNA levels, normalized against GAPDH, in PC12 cells expressing pCIneo or FH-HuD; p = 0.8661, p = 0.8767; n= 3. **(K)** Western blot images of HuD protein levels in siControl (siCon) and siHuD-transfected PC12 cells. β-Actin was used as a loading control (upper panel). Quantification of HuR mRNA levels, normalized to GAPDH, in these cells (lower panel); p = 0.0022; n = 3. Data are presented as means ± SD; ns, not significant; *p < 0.05; **p < 0.01; ***p < 0.001; ****p < 0.0001. P values were calculated using a two-tailed paired Student’s t-test. Neurites were traced in “red” in panels C and F.

Further analysis showed that a reciprocal regulation between HuD and HuR occurs in differentiating neurons. FH-HuD expression decreased HuR levels in PC12 cells (**Fig. 2H-I**), while their mRNA levels stayed the same (**Fig. 2J**). However, knocking down HuD with siRNA increased HuR mRNA levels (**Fig. 2K**). This suggests that a negative feedback mechanism involving protein stability and translation efficiency, rather than a change in transcriptional control, may be exerted by HuD or HuR on each other. Interestingly, this antagonistic interaction was not seen in non-neuronal HeLa and HEK293 cells. HuD expression does not significantly alter HuR mRNA levels in HEK293 and HeLa cells (**Fig. S2A-B**), suggesting that a cell-type-specific regulatory mechanism exists between HuR and HuD. In differentiating PC12 cells, which mimic sympathetic neurons, the neuronal ELAVL4 HuD and ELAVL1 HuR mutually repress each other.

### HuD causes neuronal differentiation by lowering miRNA activity by “sponging” and exporting repressive miRNAs from differentiating neuronal cells

Previous studies have shown that neuronal differentiation in PC12 cells is accompanied by a decline in let-7a activity, which is both sufficient and necessary to lead to the upregulation of the target gene *KRAS*, a key regulator of differentiation-related signalling pathways (16, 17). In the NGF-withdrawn condition, let-7a activity lowering is crucial for protecting against neuronal cell death (17). Consistent with these findings, we observed a significant reduction in let-7a activity upon FH-HuD expression. Luciferase reporter assays using constructs expressing reporter Renilla luciferase mRNA containing three bulged let-7a binding sites revealed decreased repression of reporter mRNA in FH-HuD-expressing cells, indicating diminished let-7a activity (**Fig. 3A-B**). This coincided with a marked increase in *KRAS* mRNA levels (**Fig. 3A, C**), supporting the idea that HuD suppresses let-7a function to upregulate differentiation-promoting genes.

**Figure 3.**
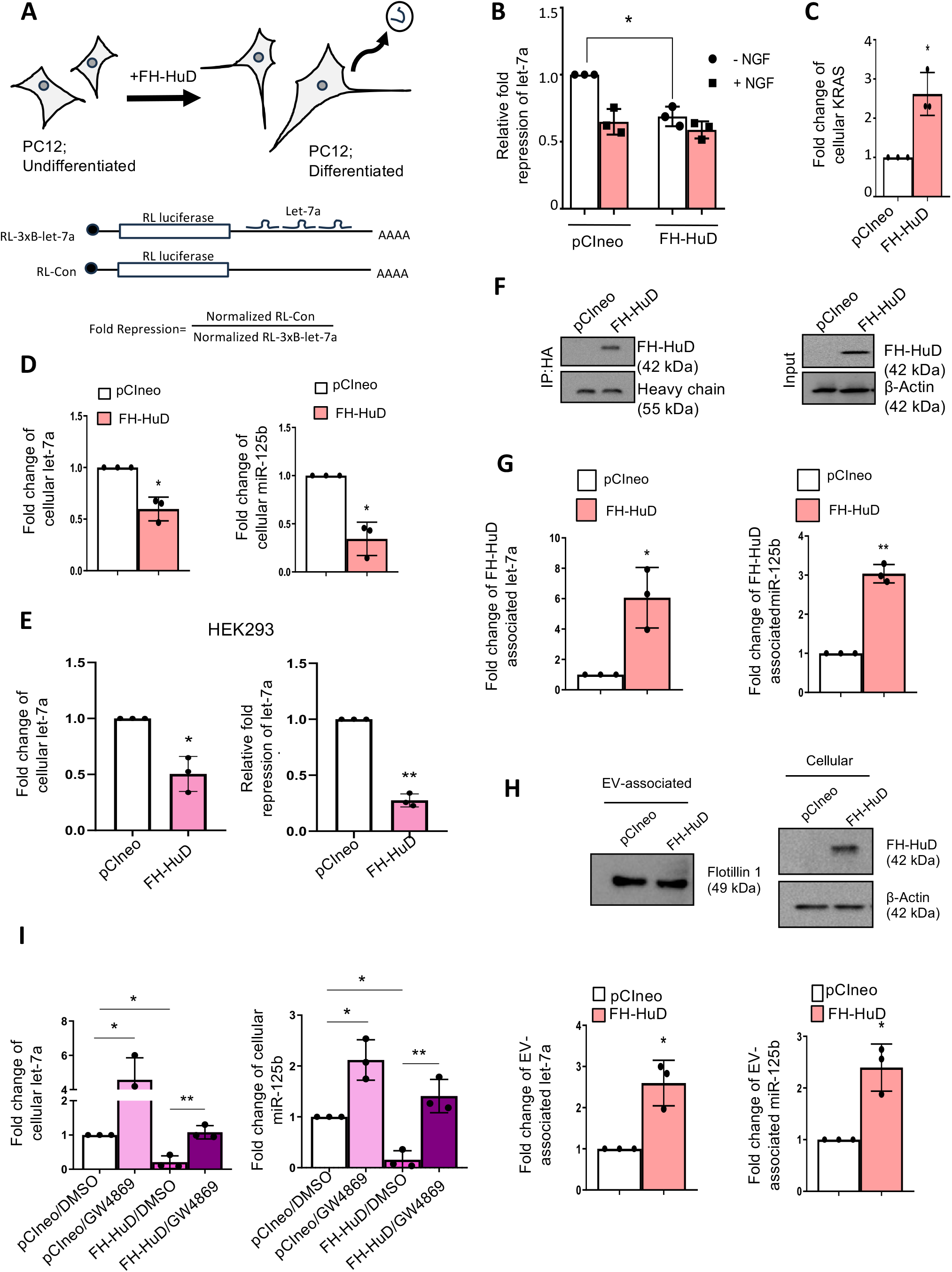
EV-mediated Export of miRNAs by HuD regulates miRNA activity in differentiating PC12 cells. **(A)** Scheme of the experiment and constructs of Luciferase reporters. **(B)** Luciferase assay showing let-7a repressive activity in pCIneo- and FH-HuD-expressing PC12 cells (with or without 72 hours of NGF treatment). p□=□0.0177; n□=□3.**(C)** qRT-PCR analysis of KRAS mRNA levels normalized to GAPDH in pCIneo- and FH-HuD-expressing PC12 cells. p□=□0.0361; n□=□3. **(D)** Cellular let-7a (left) and miR-125b (right) levels in pCIneo versus FH-HuD-expressing PC12 cells. p□=□0.0263 and p□=□0.0223, respectively; n□=□3. **(E)** qRT-PCR-based quantification of cellular let-7a miRNA levels normalized against cellular U6 levels in pCIneo- or FH-HuD-expressing HEK293 cells. p = 0.0316, n = 3 (left). Quantification of fold repression of let-7a miRNA using Luciferase assay in pCIneo- or FH-HuD-expressing HEK293 cells. p = 0.0022, n = 3 (right). **(F-G)** Immunoprecipitation of FH-HuD from pCIneo- or FH-HuD-expressing PC12 cells. Western blot was done for HA-HuD in input and IP materials (F), and qRT-PCR quantification of associated let-7a and miR-125b (G) levels normalized to pulled-down FH-HuD. p□=□0.0479 (let-7a), p□=□0.0044 (miR-125b); n□=□3. **(H)** Western blot showing Flotillin-1 (EV marker) level in EVs from pCIneo- and FH-HuD-expressing PC12 cells. EV-associated let-7a (lower left) and miR-125b (lower right) levels normalized to Flotillin-1. p□=□0.0377 and p□=□0.0339; n□=□3. **(I)** Cellular let-7a (left) and miR-125b (right) levels in pCIneo or FH-HuD-expressing PC12 cells treated with DMSO or GW4869. p□=□0.0399, 0.0176, 0.0049 (let-7a); p□=□0.0397, 0.0143, 0.0043 (miR-125b); n□=□3. Data representmeans ± SD; ns, not significant; *p < 0.05; **p < 0.01; ***p < 0.001; ****p < 0.0001. P values calculated using two-tailed paired Student’s t-test.

To determine whether this effect extended to other inhibitory miRNAs, we analysed the levels of miR-125b, another known key regulator of neuronal differentiation (18). Notably, expression of HA-HuD also led to a similar downregulation of both let-7a and miR-125b miRNAs (**Fig. 3D**). Interestingly, a comparable effect was observed in non-neuronal HEK293 cells, where FH-HuD expression led to a significant reduction in both the expression and activity of let-7a (**Fig. 3E**). Collectively, these findings suggest that HuD can modulate miRNA levels independent of cell types.

To explore the mechanism behind the reduction of miRNA activity, we observed HuD’s association with miRNAs in cells expressing FH-HuD. Analysis of RNA isolated from immunoprecipitated FH-HuD from differentiated PC12 cells and estimating associated miRNAs. HuR is known to bind to candidate miRNAs by sponging them out from the Ago2 protein and facilitating their extracellular export via EVs, thereby promoting loss of miRNA activity, as noted in hepatic and macrophage cells (10, 19). Does HuD perform similar functions in reducing miRNA activity in FH-HuD expressing cells? We isolated extracellular vesicles (EVs) from control and differentiated PC12 cells and found no significant change in EV size and a moderate decrease in EV number (**Fig. S3A**). However, the miRNA content in EVs derived from NGF-differentiated cells(+NGF) increased compared to undifferentiated cells (−NGF) (**Fig.S3A-C)**. With FH-HuD expression, we observed a similar increase in miRNA export via EVs occurring in FH-HuD-expressing PC12 cells. Extracellular vesicles were isolated from naïve or differentiated cells, as well as from cells transfected with pCIneo (control) or FH-HuD expression plasmid, and confirmed for EV-marker protein Flotillin-1 (**Fig. 3H**). qRT-PCR analysis revealed a significant increase in the level of let-7a and miR-125b in EVs following FH-HuD expression (**Fig. 3H**). These results suggest that HuD promotes the extracellular export of let-7a and miR-125b, contributing to their decreased activity in FH-HuD-expressing differentiating PC12 cells. To confirm that the decrease in cellular miRNA levels was contributed by EV-mediated export, we treated PC12 cells with GW4869, an inhibitor of exosome/EV production. In both control and FH-HuD-expressing cells, GW4869 treatment resulted in a significant accumulation of let-7a and miR-125b within the cells (**Fig. 3I**), indicating that their export depends on extracellular vesicles biogenesis, which is affected by GW4869, a chemical compound known for its ability to inhibit neutral sphingomyelinase (N-SMase) activity, an enzyme essential for Exosome biogenesis (20). HuD binds to these miRNAs and likely helps incorporate them into EVs. It plays a role in modulating PC12 cell differentiation by exporting inhibitory miRNAs via EVs, a process also involving HuR in non-neuronal or naïve PC12 cells. HuD binds let-7a and miR-125b, aiding their removal from the cytoplasm, which lifts repression on targets like KRAS. This supports a reciprocal interaction model between HuR and HuD for miRNA export, crucial for neuronal differentiation.

## Discussion

During neuronal differentiation, cells transit from proliferative progenitors to fully developed, polarized neurons with specialized structures, such as dendrites and axons (21). This process involves changes in gene regulation mediated by RNA-binding proteins and miRNAs. These proteins control mRNA stability and translation by recruiting RNA-degrading or stabilizing factors. ELAVL1, or HuR, regulates mRNA stability, translation, and responds to stress, also affecting miRNA activity in stressed hepatic cells. Neuronal homologs HuB, C, D likely have similar roles in mRNA regulation, but their specific functions and effects on miRNAs in neurons remain unclear.

Although HuR and HuD share structural features, they have opposing roles in neuronal differentiation. HuR, expressed ubiquitously, promotes cell growth, proliferation, and stress responses by stabilizing target mRNAs and preventing miRNA repression (7). Under stress, HuR moves from the nucleus to the cytoplasm, where it affects translation and exports specific miRNAs through exosomes or EVs (9, 10, 22). Conversely, HuD, primarily in neurons, increases during differentiation, supporting neuronal maturation by stabilizing mRNAs involved in synaptogenesis and neural identity (11). In this study, HuD increased while HuR decreased during PC12 cell differentiation, indicating a regulated switch essential for neuronal development.

HuD promotes differentiation by reducing inhibitory miRNAs like let-7a and miR-125b through export via exosomes or EVs, lifting repression. During neuronal differentiation, RNA export shifts with lower HuR and higher HuD. Unlike HuR, HuD is mainly cytoplasmic, facilitating miRNA export when expressed. The specific exported miRNAs and how HuD binds them for packaging into EVs are unknown. For HuR, ubiquitination releases miRNAs into endosomes, but the role of HuD’s post-translational modifications is unclear. It’s also unknown if HuD interacts with Ago-miRNPs to displace miRNAs. Investigating HuD-miRNA binding defects and their impact on neuronal differentiation, as well as the roles of ELAVL2 and ELAVL3, remains important.

### Experimental procedures

#### Cell Culture

PC12 cells were cultured in DMEM with 10% heat-inactivated horse serum, 5% FBS, and 1% Penicillin-Streptomycin. HeLa and HEK293 cells were grown in DMEM with 10% FBS and 1% Penicillin-Streptomycin. PC12 cells were differentiated with 100□ng/mL NGF in medium containing DMEM, 0.75% horse serum, and 0.25% FBS for 72 hours, then dedifferentiated with anti-NGF antibody (1:100, Sigma).

#### Cell Transfection

All cell types were transfected with Lipofectamine 2000 (Invitrogen) per the manufacturer’s instructions. For immunoprecipitation, 2□µg of plasmid DNA was transfected into cells in 60□mm dishes. For Western blotting, immunofluorescence, and RNA-FISH, 500 ng of plasmid was used in 12-well plates. Luciferase assays involved co-transfecting 20 ng of RL reporter and 200 ng of Firefly plasmids per well in 24-well plates. Transfection lasted 6 hours, then media was replaced. After 24 hours, cells were passaged to larger surfaces.

#### Western Blot

Samples mixed with 5X loading buffer and heated at 95°C for 10□minutes. Proteins were separated by SDS-PAGE, transferred to PVDF membranes, blocked with 3% BSA in 1X TBST for 1 hour, then incubated overnight at 4°C with primary antibodies. After TBST washes, membranes were treated with HRP-secondary antibodies (1:8000) for 1-1.5 hours at room temperature, washed again, and detected with chemiluminescent substrates. Imaging was done using UVP BioImager 600 and VisionWorks software (version 6.80).

#### Immunofluorescence

Cells on coverslips were fixed with 4% paraformaldehyde in PBS for 30 minutes, followed by three washes. Permeabilization and blocking used PBS with 10% goat serum, 20% BSA, and 0.1% Triton X-100 for 30 minutes. Coverslips incubated overnight at 4°C with primary antibodies in PBS with 20% BSA. After three PBS washes, secondary antibodies with Alexa Fluor were added and incubated for 1 hour at room temperature in the dark. Excess was washed away with PBS. Samples were mounted with Vectashield containing DAPI.

#### Immunoprecipitation

Cells were lysed in buffer with 20□mM Tris-HCl (pH□7.5), 150□mM KCl, 5□mM MgCl_2_, 1□mM DTT, 0.5% Triton X-100, 0.5% sodium deoxycholate, 1× PMSF, and RNase inhibitor, then incubated 30□min at 4□°C. Lysates were clarified by centrifugation at 3,000g for 15□min at 4□°C and incubated overnight with antibody-bound protein G agarose beads (Invitrogen). The next day, beads were washed three times with IP buffer (20 mM Tris-HCl [pH 7.5], 150 mM KCl, 5 mM MgCl_2_, 1 mM DTT, 1 mM PMSF, RNase inhibitor). Beads were divided for RNA and protein extraction, then analyzed by qRT-PCR and Western blot, respectively.

#### RNA Isolation and qRT-PCR

Total RNA was extracted with TRIzol or TRIzol LS (Invitrogen). mRNA and miRNA levels were quantified via two-step qRT-PCR. For mRNA, 200 ng of RNA was reverse-transcribed using the Eurogentec Reverse Transcriptase Core Kit and amplified with MESA GREEN qPCR Master Mix. For miRNA, 200□ng RNA was analysed with TaqMan miRNA-specific primers (Invitrogen), using equal volumes for FH-HuD-associated and exosomal miRNAs. Data analysis used the comparative Ct method, with GAPDH and U6 snRNA as internal controls. Western blots validated the presence of mRNA and miRNA in IP and EV samples. qPCR was performed on Bio-Rad CFX96™ and Applied Biosystems 7500. Reverse transcription: 25□°C for 10□min, 48□°C for 30□min, 95□°C for 5□min. qPCR: initial denaturation at □C for 5□min, then 40 cycles of 95□°C for 15□sec and 60□°C for 1□min.

#### Luciferase Assay

miRNA repression was assessed via dual-luciferase reporter assay. Cells were lysed with passive lysis buffer (Promega) and analyzed with the Dual-Luciferase Reporter Assay System on a VICTOR X3 plate reader (PerkinElmer). Renilla luciferase values were normalized to firefly luciferase to determine repression levels.

#### EV isolation and characterization

EVs were isolated using a modified protocol based on a previous study (23). Cells were cultured in DMEM with 10% EV-depleted horse serum, 5% EV-depleted FBS, and 1% Pen-Strep. EVs were removed from FBS by ultracentrifugation at 100,000xg for 5 hours at 4□°C. Conditioned media was centrifuged at 2,000g for 15□min and 10,000xg for 30□min, then filtered through a 0.22□μm filter. The supernatant was layered onto a 30% sucrose gradient and ultracentrifuged at 100,000g for 90□min (SW41Ti, Beckman). The EV layer was collected, washed with PBS, and pelleted again at 100,000g for 90□min. The pellet was resuspended in PBS. For characterization, EVs were diluted 1:10 in PBS and analysed with a Nanosight NS-300 for particle count and size distribution.

#### Image Capture and Post-Capture Image Processing

All confocal imaging was performed using a Zeiss LSM800 confocal microscope, and image processing and analysis were carried out using Imaris 7 software, developed by Bitplane AG. The neurite length of differentiated PC12 cells was measured in micrometers (µm) using Zen software.

#### Statistical Analysis

All graphs and statistical analyses were conducted using Prism software (v5.00 and v8.00; GraphPad, San Diego, CA). Nonparametric two-tailed unpaired and paired t-tests were used for statistical evaluation. P values less than 0.05 were considered statistically significant, whereas P values greater than 0.05 were deemed not necessary. Error bars represent the means ± standard deviation (SD).

## Data availability

All data supporting the findings of this study are available from the corresponding authors upon request.

## Supporting information

This article contains supporting information.

## Conflict of interest

The authors declare that they have no conflict of interest.

## Acknowledgements

We thank Witold Filipowicz and Gunter Meister for the constructs used in this study. We also thank the Council for Scientific and Industrial Research (CSIR) for the SR. SNB was supported by the Swarnajayanti Fellowship (DST/SJF/LSA-03/2014-15) from the Department of Science and Technology, Govt. of India. The work received support from a High-Risk High-Reward Grant (HRR/2016/000093) from the same department. SNB is supported by the Start-Up Support Grant from the University of Nebraska, USA, and K.M. acknowledges the Lieberman Research Award from the Department of Anesthesiology, UNMC.

## CRediT Author Contribution

**Sritama Ray**: Data Curation, Formal Analysis, Investigation, Methodology. **Suvendra N Bhattacharyya**: Data Curation, Formal Analysis, Investigation, Writing-Review and Editing, Conceptualization, Methodology, Funding Acquisition, Supervision. **Kamalika Mukherjee**: Data Curation, Formal Analysis, Investigation, Writing-Review and Editing, Conceptualization, Methodology, Supervision.

## Notes

### Competing Interest Statement

The authors have declared no competing interest.

