## Supplemental File for "Change of Guards: ELAVL Proteins Switch miRNA Export Responsibility to Regulate Neuronal Differentiation"

**Running Title:** miRNA Export by ELAV Proteins


**Authors**

Sritama Ray[1], Suvendra N Bhattacharyya[2], Kamalika Mukherjee[3,#]

**Affiliations**

[1]CSIR-Indian Institute of Chemical Biology, Kolkata, West Bengal, 700032, India.

[2]Department of Pharmacology and Experimental Neuroscience, University of Nebraska Medical Center, Omaha, Nebraska, 68106, USA

[3]Department of Anesthesiology, University of Nebraska Medical Center, Omaha, Nebraska, 68106, USA

### Corresponding author.

**Supplementary Figures**

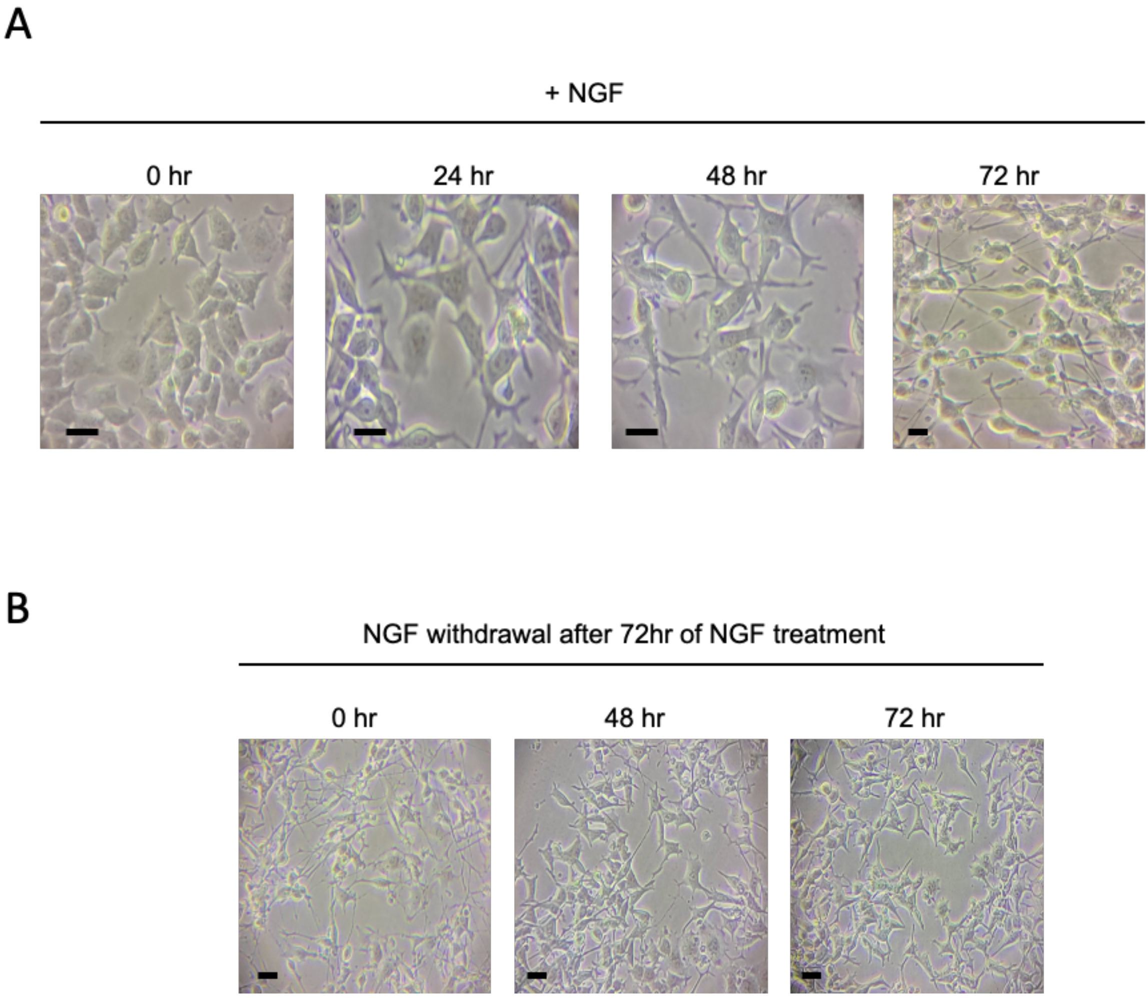


**Figure S1 PC12 differentiated with NGF and dedifferentiated without NGF in the culture medium.**
(A) Images of PC12 cells differentiated with NGF (100ng/mL) in differentiating medium for increasing time.
(B) Images of 72h NGF-differentiated PC12 cells after placing them in NGF-depleted medium for 0, 48, or 72h.
Scale Bar 10 μm.

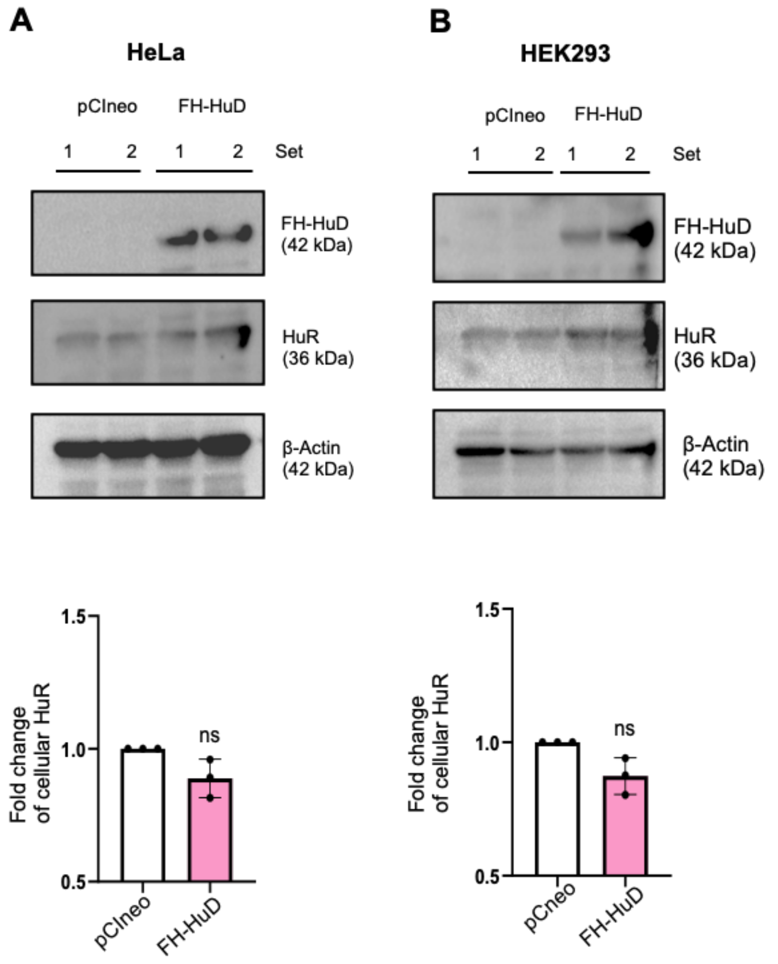


**Figure S2 Effect of FH-HuD on HuR expression in HeLa and HEK293 cells**

**(A)** Western blot images showing HuR protein levels in HeLa cells transfected with pCIneo or FH-HuD. β-Actin serves as the loading control (top). The lower panel shows qRT-PCR quantification of HuR mRNA normalized to GAPDH in the same cells; $p = 0.1163$, $n = 3$.

**(B)** Western blot images of HuR protein levels in HEK293 cells transfected with pCIneo or FH-HuD. β-Actin as the loading control (top). The bottom panel presents qRT-PCR measurements of HuR mRNA normalized to GAPDH; $p = 0.0861$, $n = 3$.

Data are expressed as means ± SD; ns, not significant; *p < 0.05; **p < 0.01; ***p < 0.001; ****p < 0.0001. p-values were calculated using a two-tailed paired Student's t-test.

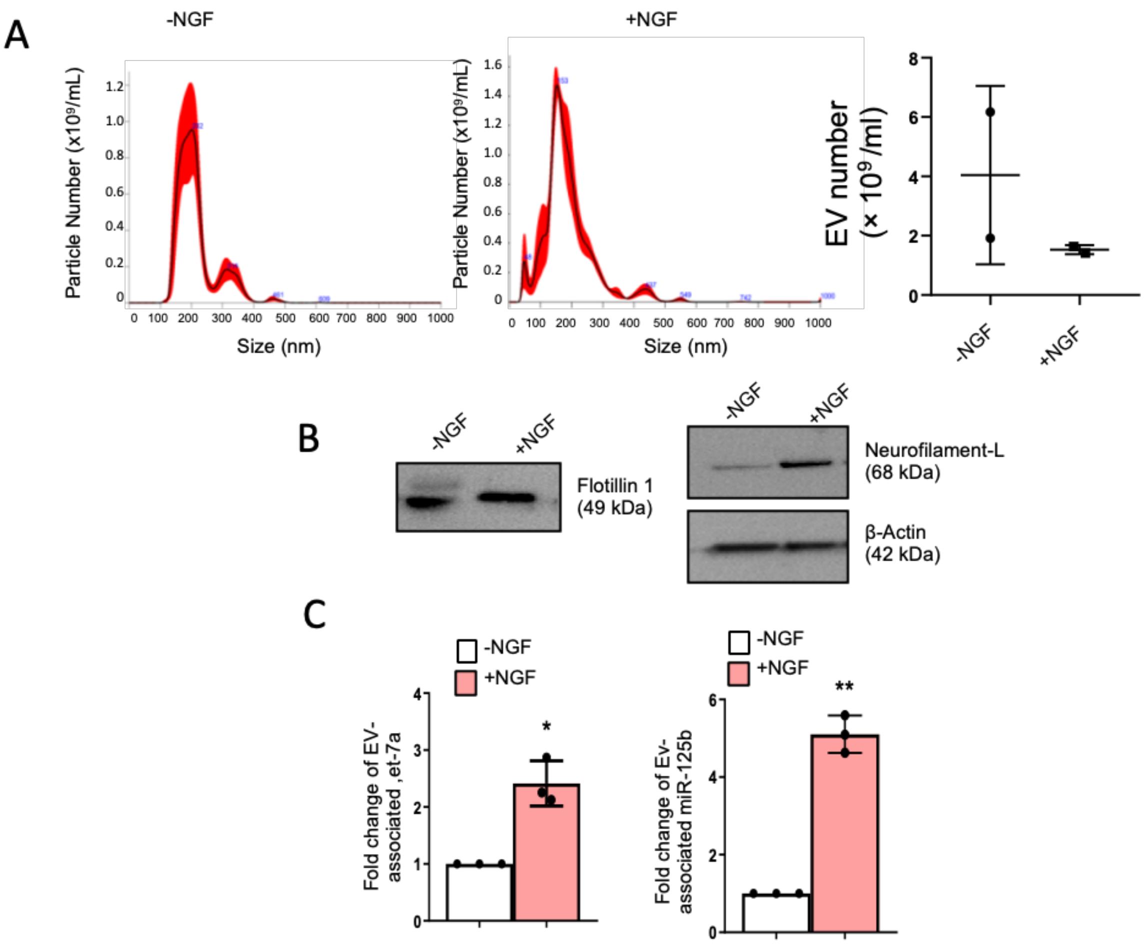


**Figure S3 Increased export of let-7a and miR-125b from differentiated PC12 cells**

**(A)** Size distribution of EVs isolated from undifferentiated vs 72 hours NGF differentiated (with 100 mg/ml) PC12 cells by NTA analysis. The right panel represents the EV concentration in culture supernatants isolated from non-differentiated versus differentiated PC12 cells.

**(B)** Western blot showing Flotillin-1 (EV-marker) level in EVs (left) and cellular Neurofilament-L from undifferentiated (-NGF) and differentiated (+NGF) PC12 cells. β-Actin was used as a loading control for cellular extracts.

**(C)** EV-associated let-7a (left) and miR-125b (right) levels normalized to Flotillin-1. $p = 0.0377$ and $p = 0.0339$; $n = 3$.

Data are expressed as means ± SD; ns, not significant; *$p < 0.05$; **$p < 0.01$. p-values were calculated using a two-tailed paired Student's t-test.

**Table S1 List of primary antibodies.**

| Antigen name | Raised in | Source | Catalogue No. | Dilution for WB | Dilution for IF/IP |
|---|---|---|---|---|---|
| HA | Rat | Roche | 11867423001 | 1:1000 | 1:100 |
| β-Actin (HRP) | Mouse | Sigma | A3854-200UL | 1:10000 | - |
| HuR | Mouse | Santa Cruz | SC 5261 | 1:1000 | 1:100 |
| HuD | Mouse | Santa Cruz | SC 28299 | 1:1000 | 1:100 |
| Neurofilament-L | Mouse | CST | #2835 | 1:1000 | - |
| Flotillin 1 | Rabbit | CST | #18634S | 1:1000 | - |

**Table S2 List of secondary antibodies.**

| Secondary Antibody | Source | Catalogue No. | Dilution |
|---|---|---|---|
| HRP Goat anti-rat | Invitrogen | 62-9520 | 1:8000 |
| HRP Goat anti-rabbit | Invitrogen | #65-6120 | 1:8000 |
| HRP Goat anti-mouse | Invitrogen | #62-6520 | 1:8000 |
| Alexa Fluor 568 (rat) | Invitrogen | A11077 | 1:500 |
| Alexa Fluor 568 (mouse) | Invitrogen | A11004 | 1:500 |

**Table S3 List of plasmids, siRNA and synthetic oligos.**

| Name | Source/Reference | Description |
|---|---|---|
| GFP-Dcp1a | Kind gift from Witold Filipowicz | Dcp1a cloned in frame with GFP |
| RL-Con | Kind gift from Witold Filipowicz | Plasmid with humanized Renilla Luciferase coding region |
| RL-3XB-let7a | Kind gift from Witold Filipowicz | Plasmid with 3 let-7a binding sites downstream of Renilla Luciferase coding region |
| pCI-neo | Promega | pCI-neo expressing vector |
| PGL3-FF | From Promega | Firefly luciferase under Simian Virus 40 (SV40) Promoter |
| FH-HuD | Prepared by cloning | Full length HuD with HA coding sequence in pIRES-Neo vector |

| | | |
|---|---|---|
| HA-HuR | Kind gift of Witold Filipowicz | Full length HuR with HA coding sequence in pCI-Neo vector |
| siHuD (rat) | Dharmacon | Catalogue No. L-108537-01-0005 |

**Table S4 List of mRNA real-time primers.**

| Target | Primer | Sequence |
|---|---|---|
| GAP43 | Forward | 5' ACGAGAAGAAGGGTGATGCA 3' |
| | Reverse | 5' CTTCGCCCTTCTTCTCCTCA 3' |
| Neurofilament-M | Forward | 5' AACGTCAAGATGGCTCTGGA 3' |
| | Reverse | 5' GTGTTGGACCTTGAGCTTGG 3' |
| HuD | Forward | 5' TCACCATTGACGGGATGACA 3' |
| | Reverse | 5' ACCTTGACGTTGTTCACTGC 3' |
| HuR | Forward | 5' TCAACTCCAGGGTCCTTGTG 3' |
| | Reverse | 5' GTTCTGGTTGGGATTGGCTG 3' |
| GAPDH | Forward | 5' CAGGGGGGAGCCAAAAGGG 3' |
| | Reverse | 5' CTTGGCCAGGGGTGCTAAGC 3' |
| KRAS | Forward | 5' AACTGGGGAGGGCTTTCTTT 3' |
| | Reverse | 5' TGTCTTGTCTTCGCTGAGGT 3' |

**Table S5 List of miRNA assays used for Taqman based quantification.**

| miRNA | Assay ID |
|---|---|
| Human let-7a (hsa-let-7a) | 000377 |
| Human miR-125b (hsa-miR-125b) | 000449 |
